# PathoFact: A pipeline for the prediction of virulence factors and antimicrobial resistance genes in metagenomic data

**DOI:** 10.1101/2020.03.24.006148

**Authors:** Laura de Nies, Sara Lopes, Anna Heintz-Buschart, Cedric Christian Laczny, Patrick May, Paul Wilmes

## Abstract

**Background:** Pathogenic microorganisms cause disease by invading, colonizing and damaging their host. Virulence factors including bacterial toxins contribute to their pathogenicity. Additionally, antimicrobial resistance genes allow pathogens to evade otherwise curative treatments. To understand causal relationships between microbiome compositions, functioning, and disease, it is therefore essential to identify virulence factors and antimicrobial resistance genes in metagenomic datasets. At present, there is a clear lack of computational approaches to simultaneously identifying these factors. Here we present PathoFact, a tool for the contextualized prediction of virulence factors and antimicrobial resistance genes in metagenomic data.

**Results:** PathoFact predicts virulence factors, bacterial toxins and antimicrobial resistance genes with high accuracy (0.92, 0.83 and 0.99) and specificity (0.96, 0.99 and 0.98), respectively. The performance of PathoFact was furthermore demonstrated on three publicly available case-control metagenomic datasets representing an actual infection as well as chronic diseases in which either pathogenic potential or bacterial toxins were predicted to play a role. With PathoFact, we identified virulence factors (including toxins) and antimicrobial resistance genes, and identified signature genes which differentiated between the disease and control groups.

**Conclusion:** PathoFact is an easy-to-use, modular, and reproducible pipeline for the identification of virulence factors, toxins and antimicrobial resistance genes in metagenomic data. Additionally, PathoFact combines the prediction of these pathogenicity factors with the identification of mobile genetic elements. This provides further depth to the analysis by considering the genomic context of the pertinent genes. Furthermore, each module (virulence factors, toxin and antimicrobial resistance genes) of PathoFact is also a standalone component, making it a flexible and versatile tool. PathoFact is freely available online at https://git-r3lab.uni.lu/laura.denies/PathoFact.

## Introduction

Most of the microorganisms constituting the human microbiome are commensals. They contribute essential functions to the human host and contribute to its physiological development. In contrast, pathogenic microorganisms including bacteria, viruses, fungi, and protozoa cause disease by invading, colonizing and damaging the host. Virulence factors, including bacterial toxins contribute to this pathogenicity by enhancing not only the infectivity of pathogenic bacteria but also by exacerbating antimicrobial resistance which in turn restricts treatment options [1].

Virulence factors enable in particular pathogenic microorganisms to colonize host niches ultimately resulting in tissue damage as well as local and systemic inflammation. They include a wide range of factors which contribute both directly and indirectly to disease processes, and which are important for pathogens to establish an infection [2]. These virulence traits include cell-surface structures, secretion machineries, siderophores, regulators, etc. [3,4]. However, of all virulence factors employed by pathogens, bacterial toxins count as the most important effector proteins. Different types of bacterial toxins have evolved over time to counteract human defenses. These can be roughly categorized into two groups: the cell-associated endotoxins and the extracellular diffusible exotoxins. Exotoxins are typically polypeptides and proteins that act to stimulate a variety of host responses either through direct action with cell receptors or via enzymatic modulation [5,6].

Partly through the utilization of these virulence factors, and toxins in particular, pathogenic microorganisms have been a major cause of infectious diseases including in the context of viral co-infections. The development and medical use of antibiotics has limited the development and spread of these pathogens by providing an effective treatment for bacterial infections. However, the over- and mis-use of antibiotics has resulted in a global increase in antimicrobial resistance (AMR) which now threatens human health through the emergence and spread of multidrug resistant bacteria [1,7]. As a result, many pathogenic bacteria have now acquired resistance against the main classes of antibiotics which has led to a dramatic rise in untreatable infections, even resulting in the emergence of so-called “superbugs” that are extremely difficult to treat [8]. Consequently, AMR is an urgent and growing threat to public health with an estimated number of deaths exceeding ten million annually by 2050 [9,10].

The acquisition of antimicrobial resistance genes (ARGs) is not restricted to a single strain or species of bacteria. While commensal bacteria provide a source of ARGs, antimicrobial resistance can be transferred to pathogenic species through horizontal gene transfer, either through conjugation or transduction [11–13]. Therefore, to understand the emergence and spread of ARGs, it is necessary to monitor microbial communities in their native habitats. Through providing unbiased views of the genomic complements of individual microbial populations, metagenomics allows such surveys.

Compared to virulence factors, AMR has evolved on a different time scale [7]. Pathogenic microorganisms have modified and adapted their virulence to the host defense system over millions of years. Meanwhile, with an increase in selective pressure through the use of antibiotics an excessive increase in the spread and evolution of AMR has been observed in the last fifty years. However, despite the differences in evolutionary timescales and paths, virulence factors and AMR share common characteristics. Most importantly, virulence factors and AMR are necessary for pathogenic bacteria to adapt to, and survive in, competitive microbial environments [7]. Additionally, both virulence and resistance mechanisms are mostly transferred between bacteria by horizontal gene transfer [14]. Furthermore, both processes make use of similar systems that activate or repress the expression of various genes, including efflux pumps, porins, cell wall alterations, and two-component systems [15–17]. Therefore, although AMR in itself is not a virulence factor, in environments with selective antibiotic pressure, opportunistic pathogens are able to colonize through acquisition or presence of AMR [1].

Considering the burden of bacterial infections in which virulence factors and ARGs play crucial functions, it is important to be able to identify these in microbial communities. The advent of high-throughput DNA sequencing provides a powerful means to profile the full complement of DNA derived from genomic extracts obtained from a wide range of environments [11]. However, currently there is a clear lack of computational approaches to simultaneously identifying these different factors in metagenomic datasets. Various tools exist for the prediction of ARGs themselves, such as deepARG [11], ResFinder [18] and ARGsOAP [19], with a very few prediction tools for virulence factors existing as well, such as MP3 [20] and VirulentPred [21]. Even though a few tools exist for the prediction of virulence factors, most are based on older databases of virulence factors which have since been expanded greatly. Moreover, there is a lack of bioinformatics tools for the specific prediction of bacterial toxin genes in particular. Furthermore, although various AMR prediction tools exist, these primarily focus on the prediction of genes without differentiating between mobile genetic elements (MGEs) or on the bacterial genomes. Since MGEs are the main mechanism by which ARGs are transmitted, it is crucial to identify any relationships between ARGs and MGEs. Consequently, it is of interest to create new tools for the prediction of both virulence factors, and toxins in particular, and combine these with the prediction of ARGs and MGEs. Therefore, we have developed PathoFact, a pipeline for the prediction of virulence factors, bacterial toxins and antimicrobial resistance genes in their respective genomic contexts.

## Implementation

PathoFact is a command-line tool for UNIX-based systems that integrates three distinct workflows for the prediction of (i) toxins, (ii) virulence factors, and (iii) antimicrobial resistance genes from metagenomic data (Fig. 1a). Each workflow can be applied individually or in combination with the other workflows. Our pipeline is written in Python (version 3.6) using the Snakemake (version 5.5.4) workflow management software [22]. This implementation offers several advantages, including workflow assembly, parallelism, and the ability to resume processing following an interruption. Each step of the pipeline is implemented as a rule in the Snakemake framework specifying the input needed and the output files generated. Additionally, several rules in the workflows make use of conda environments in which necessary dependencies of the rule are stored. Moreover, the use of conda environments makes it possible to incorporate prediction tools dependent on older Python versions incompatible with version 5.5 of Snakemake. As such, Python, as well as a working snakemake and (mini)conda (version 4.7) [23] installation are required. PathoFact is available under https://git-r3lab.uni.lu/laura.denies/PathoFact.

**Figure 1.**
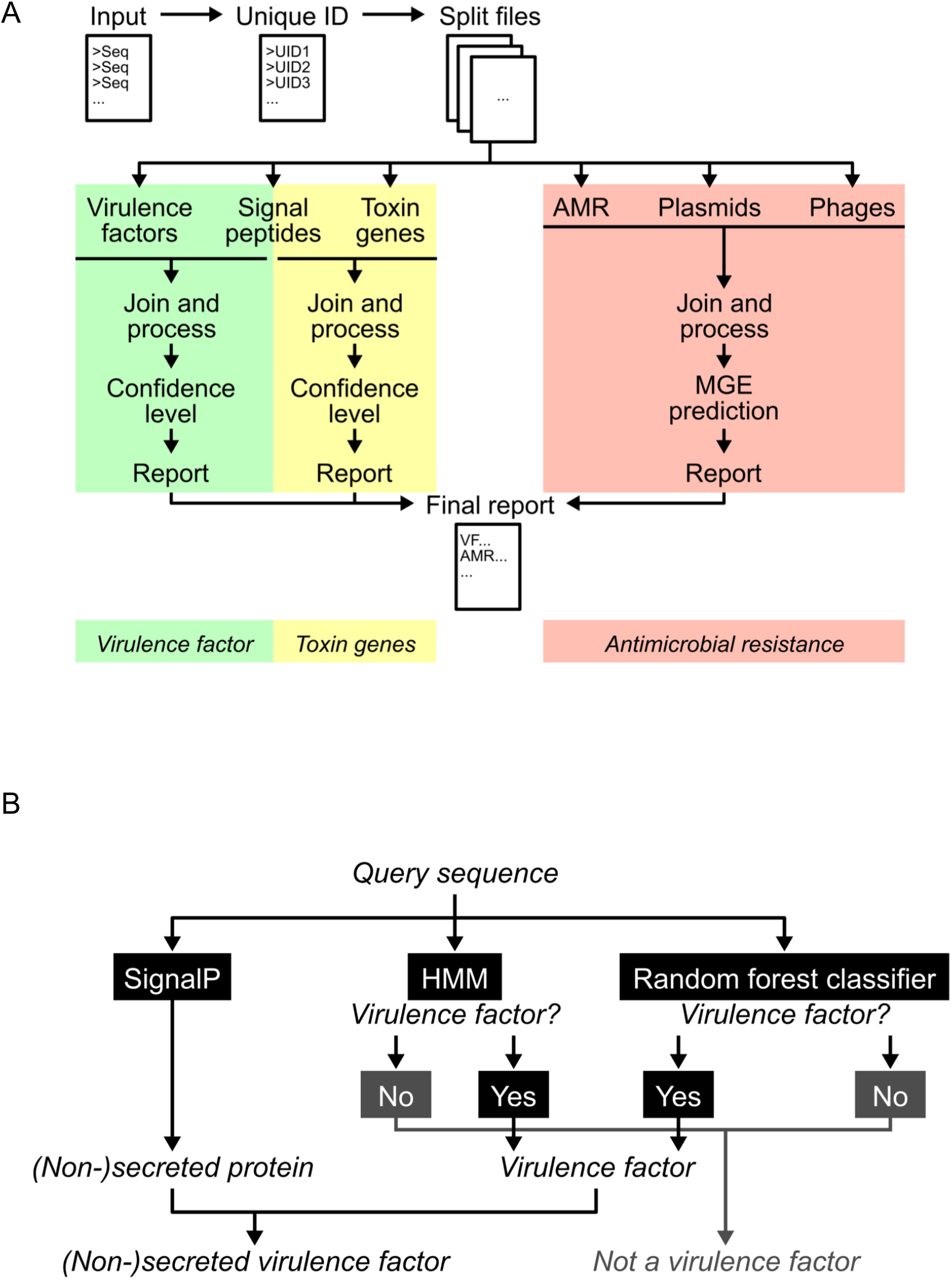
The PathoFact pipeline. **a**. Framework of the PathoFact pipeline. The pipeline consists of three different modules related to (i) virulence factors, incl. (ii) bacterial toxins and (iii) antimicrobial resistance genes. SignalP is incorporated for the prediction of secreted toxins and virulence factors. All modules can either be run independent or as one single pipeline. **b.** Classification framework for the prediction of virulence factors. The prediction of virulence factors depends on two different aspects: (i) a HMM domain database, (ii) a random forest classifier. Sequences predicted positive from both are classified as virulence factors. The incorporation of SignalP in the framework incorporates information regarding the likely secretion of the virulence factors.

The input to the PathoFact pipeline consists of: (i) an amino acid fasta file of translated gene sequences for the prediction of virulence factors, bacterial toxins and antimicrobial resistance genes, (ii) a fasta file containing nucleotide sequences of the corresponding contigs for the prediction of MGEs and resolution of genomic context, and (iii) a table consisting of a first column of contig names with the corresponding gene names in the second column to combine predictions. Contig names need to correspond to the original names provided in the fasta headers.

As a first preprocessing step, a unique identification number is given to each sequence in the fasta files to allow the merging of outputs of the different tools. Input files can be automatically split into individual subsets to promote parallelization of the workflow. This allows for faster run times and for the pipeline to process larger datasets, as is often the case with metagenomic data.

### Workflow for the prediction of virulence factors

For the prediction of virulence factors, we created a prediction tool consisting of two parts; (i) a database consisting of virulence factor HMM profiles, and (ii) a machine learning model. Hits against the virulence factor HMM database are then combined with the classification of the machine learning model to result in the final prediction (Fig. 1 b). The development of the tool was based on the MP3 software tool for the prediction of virulence factors [20]. In addition, analyses were combined with the prediction of signal peptides by SignalP [24] to distinguish between secreted and non-secreted virulence factors.

### Dataset for the predictions of virulence factors

A dataset, consisting of both a positive and negative subset, was constructed for the training of the virulence factor prediction tool. The positive subset consisted of known virulence factor sequences retrieved from the Virulence Factors Database (VFDB)[3]. All sequences were obtained from the VFDB core dataset containing (translated) gene sequences associated with experimentally verified virulence factors. The negative subset of the training set consisted of protein sequences that were retrieved from the Database of Essential Genes (DEG) [24] and which were known not to be virulence factors. For both subsets, all sequences were clustered with CD-HIT and sequences with a 90% sequence identity were removed. The resulting training set was used for (i) the training of the machine learning model, and (ii) the implementation of the HMM profiles.

### Machine learning model for the prediction of virulence factors

As part of the prediction of virulence factors, a random forest model was created [25]. A random forest model operates from decision trees and outputs classification of the individual trees while correcting for overfitting of the training set. For training of the random forest model the following five features were selected and implemented: amino acid composition (AAC), dipeptide composition (DPC), composition (CTDC), transition (CTDT) and distribution (CTDD) [26]. A feature matrix was built with rows corresponding to the sequence composition of the features. The random forest model was implemented using the pandas, numpy, pickle and scikit-learn python packages and consisted of 1600 trees with a maximum depth of 340.

### Construction of the virulence factor HMM Database

As the second part for the prediction of virulence factors, HMMs were used to construct the virulence HMM database. The corresponding HMM profiles to the training set were identified using five different databases: PFAM-A [27], TIGR [28], KEGG [29], MetaCyc [30] and Swissprot [31] [32]. Identified HMM domains were selected based on their best hit in all databases as determined by the highest bit-score obtained. HMM profiles were subsequently retrieved and the databases were concatenated to form the virulence HMM database. Binary compressed data files were constructed with the *hmmpress* function. During the prediction of virulence factors by the virulence HMM database identified HMM profiles are then separated by those originating to the positive or negative subset of the training set as well as HMM profiles ambiguous for both positive and negative subset.

### Workflow for the prediction of toxin genes

For the prediction of toxin genes, a workflow consisting of a toxin HMM database combined with SignalP version 4.0 [33] was developed. The toxin HMM database consists of bacterial toxin domains to identify toxin-related domains in the query sequences. Using the *hmmsearch* function of the HMMER3 program [34] the query sequences are searched against the collection of profiles present in the toxin HMM database. In addition, analyses are combined with SignalP [24] to differentiate between secreted and non-secreted toxins.

### Construction of the toxin HMM database

An HMM model based on a training set of known toxins was developed and implemented using the HMMER3 software. The training set was compiled from the Toxin and Toxin Target Database (T3DB)[35] with the training set derived from the DBETH prediction tool [5]. Protein sequences from the training set with a similarity greater than 90% were clustered and removed with CD-HIT-2D [36]. In a similar method to the construction of the virulence HMM database, the corresponding HMM profiles were identified from the five previously described databases. The datasets were extended with HMM profiles identified from an additional manual search of annotated bacterial toxins in the five databases. Finally, in order to have a short description of all HMM profiles present in the toxin HMM database, a toxin library was created. This lists (i) all HMM profiles, (ii) their names, (iii) their alternative names, and (iv) the original database from which the HMM profile was derived.

### Workflow for the prediction of antimicrobial resistance genes

For the prediction of ARGs, the workflow is separated into two parts; (i) the prediction of ARGs, and (ii) the prediction of MGEs. For the prediction of ARGs, the tool DeepARG was used [11]. DeepARG uses a deep learning approach that improves classification accuracy while at the same time reducing false negatives. It offers a powerful approach for metagenomic profiling of ARGs as it expands on the available databases for ARGs by combining the widely used CARD [37], ARDB [38], and UNIPROT [39] databases.

### MGEs: plasmids and phages

The prediction of MGEs is split into two parts focusing on the prediction of (i) plasmids, and (ii) phages. For the prediction of plasmids, PlasFlow [40] is used, while for the prediction of phages two tools are used (i) VirSorter [41] and (ii) VirFinder [42]. The predictions of these different tools are merged with the prediction of ARGs, to provide a final localization of the resistance genes to either MGEs or genomes. Considering the different predictions of MGEs, the final classification includes plasmid, phage, genome, unclassified, and ambiguous when localization predictions contradict each other, for example predicted to be both phage and plasmid.

### Evaluation of the PathoFact pipeline

To evaluate the performance of the PathoFact pipeline, validations were conducted for the prediction of toxins, virulence factors and AMR genes respectively (table 1). The prediction quality was evaluated by sensitivity, specificity and accuracy criteria as defined below.

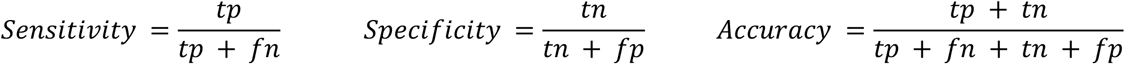

**Table 1.** Validation of the PathoFact pipeline. Evaluated performance of PathoFact regarding the prediction of virulence factors, incl. bacterial toxins, and antimicrobial resistance genes.

|  | Toxin prediction | Virulence factor prediction | DeepARG |
| --- | --- | --- | --- |
| Sensitivity | 0.777 | 0.886 | 0.988 |
| Specificity | 0.989 | 0.957 | 0.983 |
| Accuracy | 0.832 | 0.921 | 0.986 |

Where *tp* represents true positives (i.e. virulence factors (incl. bacterial toxins) or AMR gene is predicted correctly), *tn* (i.e. a gene is correctly predicted not to be a virulence factor, toxin genes or AMR gene), *fp* false positive (i.e., a gene incorrectly identified as a virulence factor, toxin genes or AMR gene), and *fn* false negatives (i.e, a virulence factor, toxin genes or AMR gene is incorrectly identified as such).

### Validation of virulence factors

A validation dataset was constructed to assess the unbiased performance of the prediction of virulence factors. As done for the training set, the validation set consisted of a positive subset of 2639 sequences (VFDB database) and a negative subset of 2628 (DEG database) sequences. Importantly, the sequences in the validation dataset were removed from the training set to reduce overfitting.

### Validation of toxin genes

For the validation of toxin genes, a validation dataset containing both positive and negative subsets was constructed. The positive subset was constructed from a manual search in the EMBL-EBI database for bacterial toxins. The results were limited to protein sequences described in the UniProtDB. Further filtering of the protein sequences removed sequences with uncertain predictions (i.e. hypothetical, probable). To limit redundancy within the dataset, sequences were clustered in terms of similarity by using a 90 % identity cut-off. Furthermore, to limit redundancy within the training set, sequences with a similarity of greater than 90% were discarded. The remaining 202 positive sequences were combined with 202 selected sequences from the negative dataset, consisting of housekeeping genes to result in the validation dataset.

### Validation of AMR prediction

For the prediction of AMR genes, the deepARG prediction tool was used. DeepARG has proven to be more accurate than most AMR prediction tools with a great reduction in false negatives [11]. Before including DeepARG in the pipeline, the prediction tool was tested using the NCBI’s resistance gene database [43]. The validation set consisted of a positive subset (5265 sequences) combined with a negative subset of equal size.

### Data analysis and data availability

Metagenomic sequences were obtained from the European Bioinformatics Institute-Sequence Read Archive database, with accession numbers PRJNA297269 (Milani *et al.)*, PRJNA281366 (Tett *et al.)* and ERP019674 (Bedarf *et al.)*. Metagenomic reads were processed and assembled using IMP [44]. The resulting fasta files containing the assembled contigs and genes were used as input for PathoFact. For analyses of the predictions, FeatureCounts [45] was used to extract the number of reads per functional category. Thereafter, the relative abundance of the toxin genes was calculated using the Rnum_Gi method described by Hu *et al[46]*. Additionally, the R DESeq2 [47] package was used to analyze the differential abundance of virulence factors, toxins and AMR genes.

## Results and Discussion

### Validation of the PathoFact pipeline

For the prediction of virulence factors an overall sensitivity of 0.886, specificity of 0.957 and an accuracy of 0.921 were achieved (Table 1). A high specificity means that only few false negatives were predicted. Additionally, the tool was compared to the MP3 tool for the prediction of virulence factors (Supplementary Table 1). With an overall sensitivity of 0.886, specificity of 0.957 and accuracy of 0.921, PathoFact scored overall better than MP3 which scored 0.125, 0.992, 0.558 respectively. In addition to the prediction of virulence factors, for the prediction of bacterial toxins itself an overall sensitivity of 0.777, specificity of 0.989 and accuracy of 0.832 were obtained.

Finally, for the prediction of ARGs an overall sensitivity of 0.988, specificity of 0.983 and accuracy of 0.986 demonstrates the precision of the deep learning approach for the prediction of AMR genes, as has also been demonstrated previously by Arango-Argoty *et al*. [11]

### Performance of PathoFact on independent metagenomic datasets

Virulence factors and toxins may contribute to dysbiosis of the microbiome and favor a pro-inflammatory environment [48]. In addition, in particular pathogenic bacteria may adapt to, and survive in, the presence of antimicrobials through acquisition or expression of AMR. Thereby, virulence factors, toxins and AMR may all contribute to the pathogenic potential of the microbiome, which in turn may have an effect on the onset and development of disease and infection. The performance of PathoFact was demonstrated using three publicly available casecontrol metagenomic datasets which were chosen considering the following criteria: representing an actual infection or a chronic disease in which either pathogenic potential or toxins are believed to play a role. The Milani *et al. [49]* study represents actual infections with *Clostridioides difficile* (CDI) in the human gut microbiome of five patients along with five healthy controls. Furthermore, skin metagenomes of five psoriasis patients along with five healthy controls from Tett *et al. [50]* were chosen to represent a chronic disease in which a pathogenic potential is believed to have a function. Additionally, from Bedarf *et al*. [51] the metagenomes of fecal microbiome of 17 early stage Parkinson’s disease (PD) patients, as well as 15 age-matched controls, was obtained to represent chronic diseases in where bacterial toxins are believed to be involved.

### Prediction of virulence factors and bacterial toxins

The predictions from PathoFact resulted in the identification of virulence factors in all three casecontrol metagenomic datasets. Furthermore, predicted virulence factors were characterized as secreted and non-secreted through the incorporation of SignalP in the pipeline. In neither of the three datasets a significant difference was found in the relative abundance of the different virulence factors between the diseased state and control (Fig. 2). Although a higher abundance was linked to CDI and a wider spread between samples can be observed in PD.

**Figure 2.**
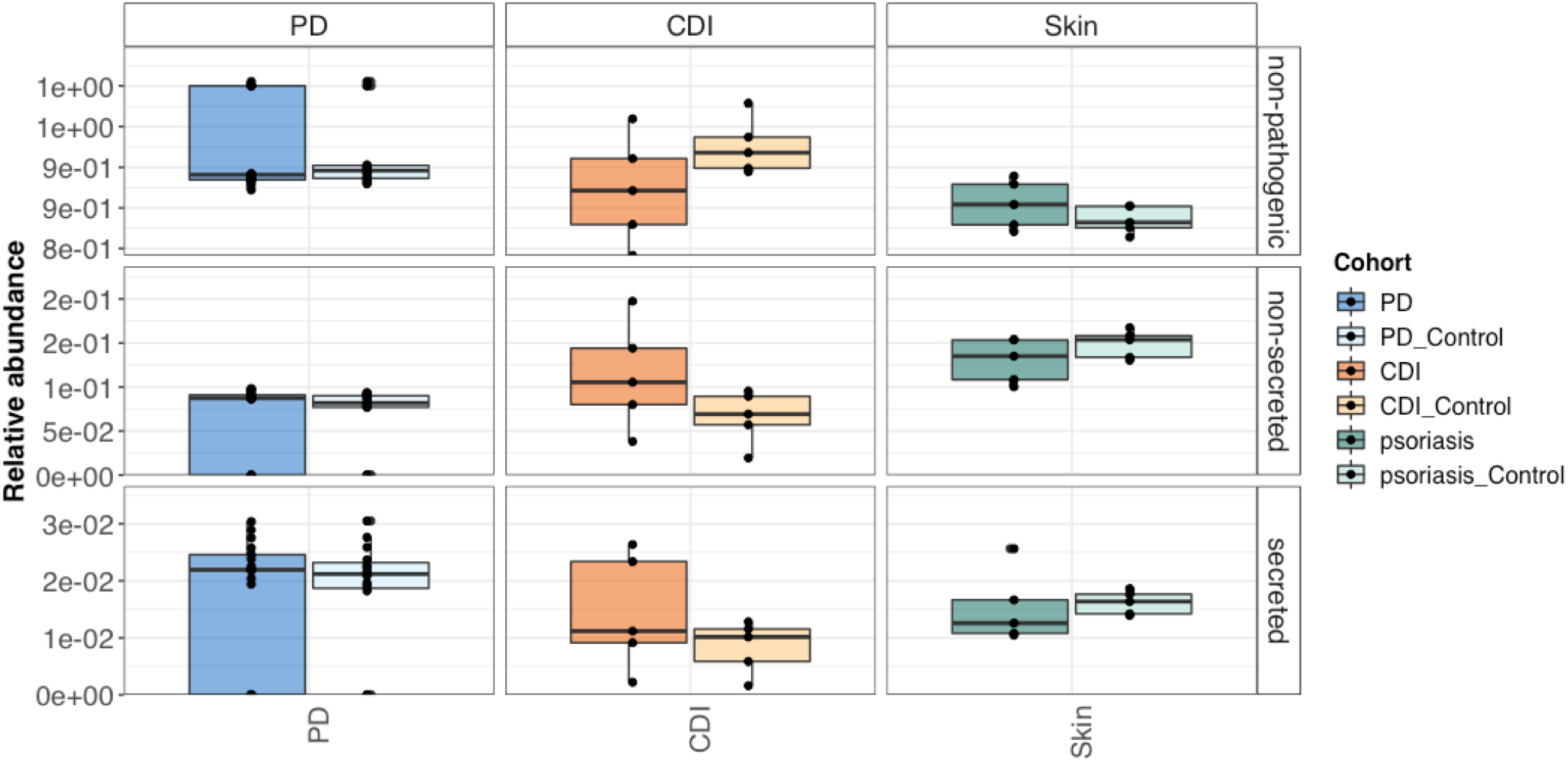
Virulence factors in diseased versus control individuals in three case-control metagenomic datasets. The relative abundances (%) of both secreted and non-secreted virulence factors as well as non-pathogenic sequences are provided.

In addition to the general prediction of virulence factors using PathoFact we identified bacterial toxins, as well as their corresponding HMM domain by which they were identified, in all three datasets. Furthermore, predicted toxin genes were characterized as secreted and non-secreted through the incorporation of SignalP in the pipeline. Both secreted and non-secreted toxins were identified in both diseased and control groups in all datasets (Fig. 3A). Moreover, we found that several bacterial toxins were differentially abundant between diseased and control (Supplementary Table 2). In the metagenomic study representing an actual infection of *Clostridioides difficile* three distinct toxin domains, PF13953, PF13954 and PF06609, were identified to be differential abundant in CDI over control (Fig. 3b). Interestingly, none of these toxin domains have yet been reported to be linked to CDI and therefore may be of interest for further research. Regarding the skin metagenomes we found a number of bacterial toxins differential abundant between cohorts (Fig. 3c). Four distinct toxin domains (K12340, PF13935, PF14449 and K11052) were found to be significantly abundant in in psoriasis over controls. Of these toxin domains only K12340 was previously linked to psoriasis [52]. Finally, regarding the PD study we found several differentially abundant bacterial toxins differential abundant between PD and control (Fig. 3d). Of these bacterial toxins two containing the PF09156 and PF09599 domains were found to be more abundant in PD. Of these domains PF09599 is among others found in invasin proteins in *Salmonella typhimurium* which has been hypothesized to be involved in Parkinson’s disease [53]. Interestingly, in all three datasets additional toxin domains were identified unknown to be linked to the diseases, therefore these domains may be of interest for further research.

**Figure 3.**
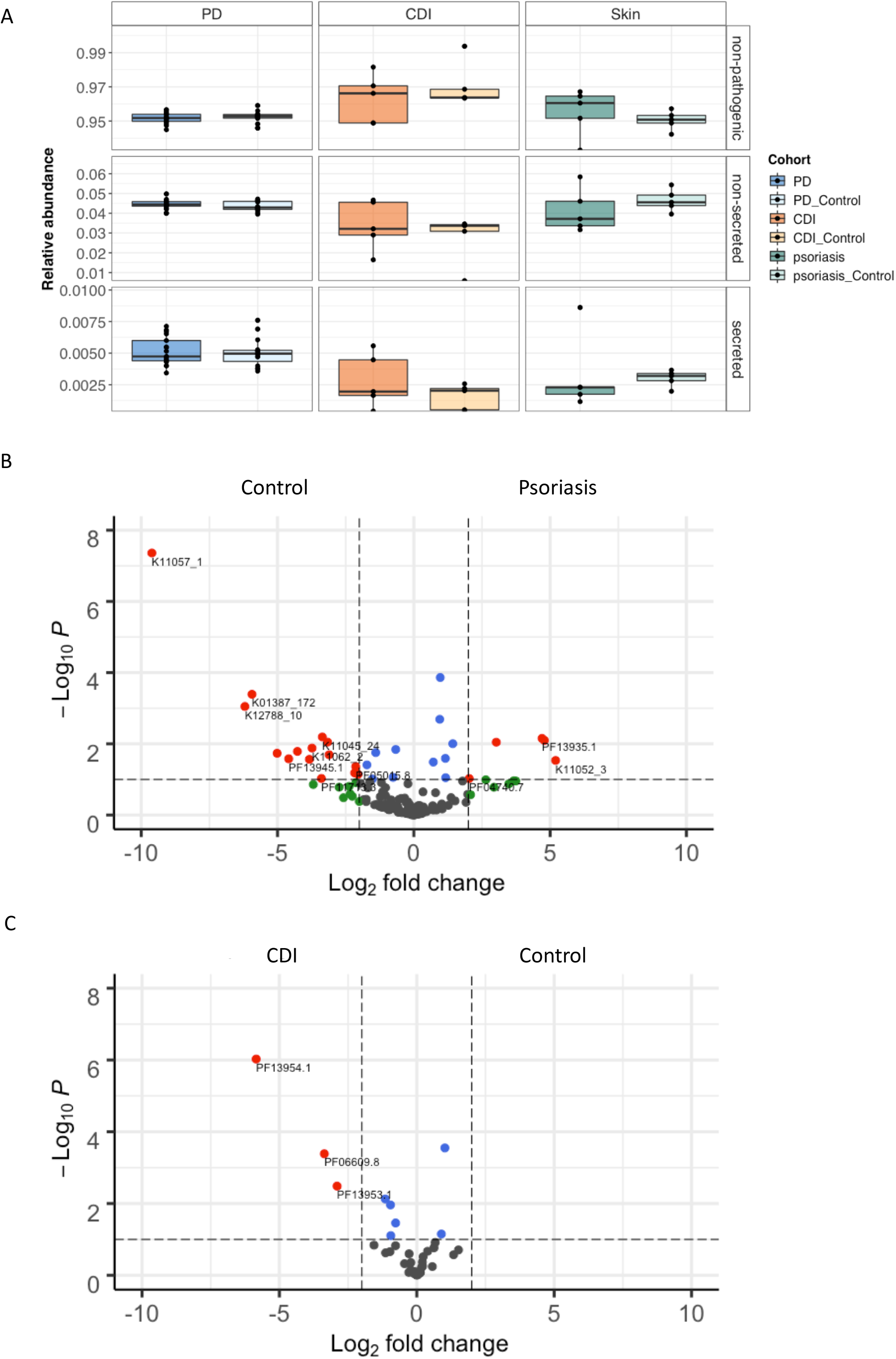
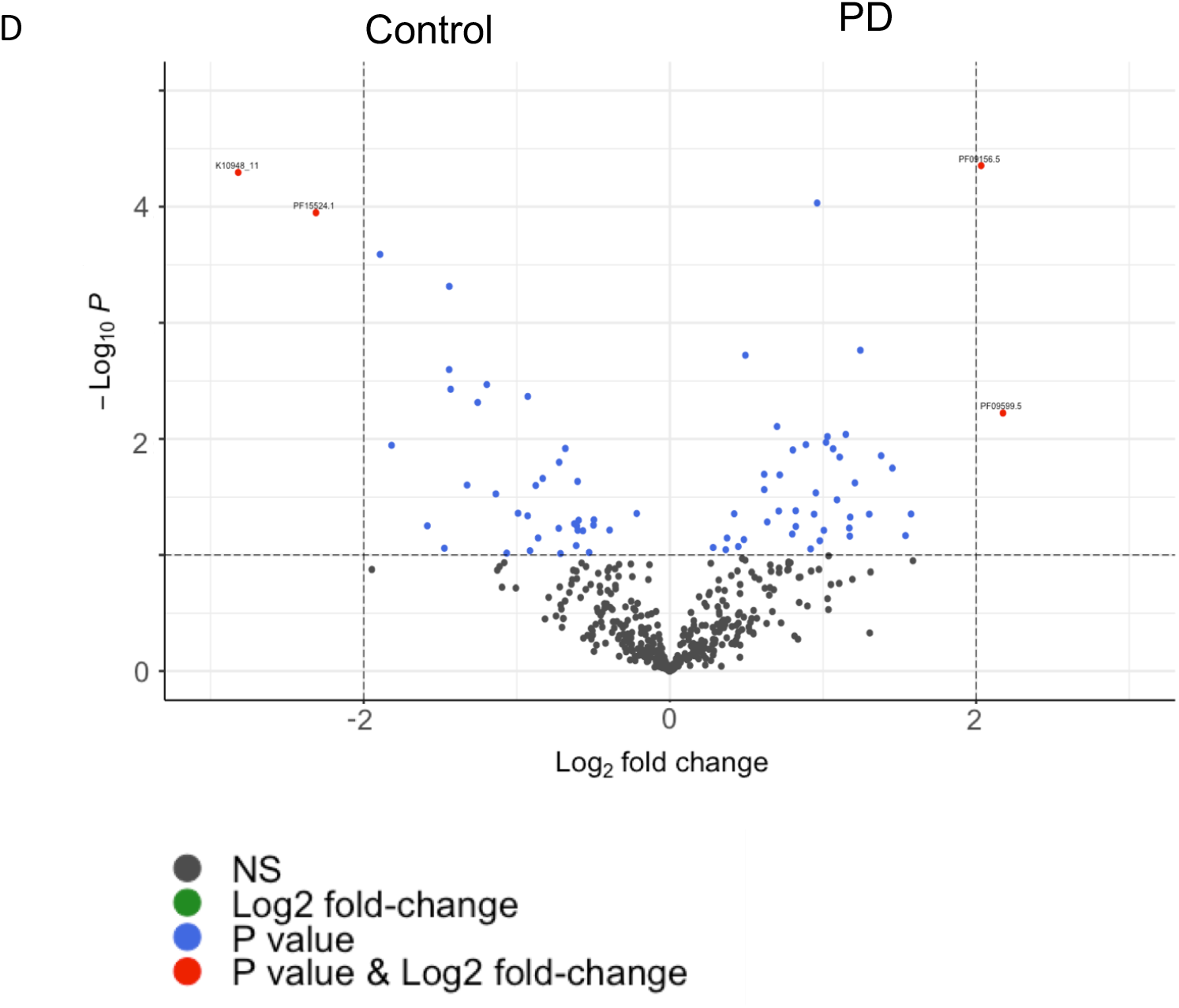
Bacterial toxins in three case-control metagenomic datasets. Bacterial toxins in disease versus control datasets **a.** The relative abundance (%) of both secreted and non-secreted bacterial toxins in diseased versus control subjects. **b.** Volcano plot depicting differentially abundant bacterial toxins in *Clostridioides difficile* infections versus control. **c.** Volcano plot depicting differentially abundant bacterial toxins in Psoriasis versus control. **d.** Volcano plot depicting differentially abundant bacterial toxins in Parkinson’s disease versus control.

### Prediction of antimicrobial resistance

Using the PathoFact pipeline we predicted the presence of antimicrobial resistance genes in all three case-control metagenomic datasets. Within the CDI datasets 25 categories of ARGs were identified (Supplementary Figure 1 a) of which six were significantly higher abundant in individuals with CDI over control (Fig. 4a). All of the identified resistance categories, i.e. aminoglycosides, fluoroquinolone, peptide and multidrug resistance, have previously been found to be associated with CDI infections [54]. In the metagenomic data of the skin microbiome 24 categories of ARGs (Supplementary Figure 1b). Interestingly, of the identified resistance categories two categories (fosfomycin and antibacterial peptides) were found to be significantly decreased within the psoriasis patients compared to the control group (Fig. 4b). Within the PD study 26 ARG categories were identified (Supplementary Figure 2c) with five categories (i.e. mupirocin, nucleoside, oxazolidinone and polyamine:peptide) significantly abundant in PD over controls (Fig. 4c). The link between antimicrobial resistance and Parkinson’s disease has been mostly unexplored thus far. However, a recently published study by Mertsalmi *et al [55]* suggests a role for antibiotics through the influence on the gut microbiome could be a factor in PD. Our present results support these initial observations.

**Figure 4.**
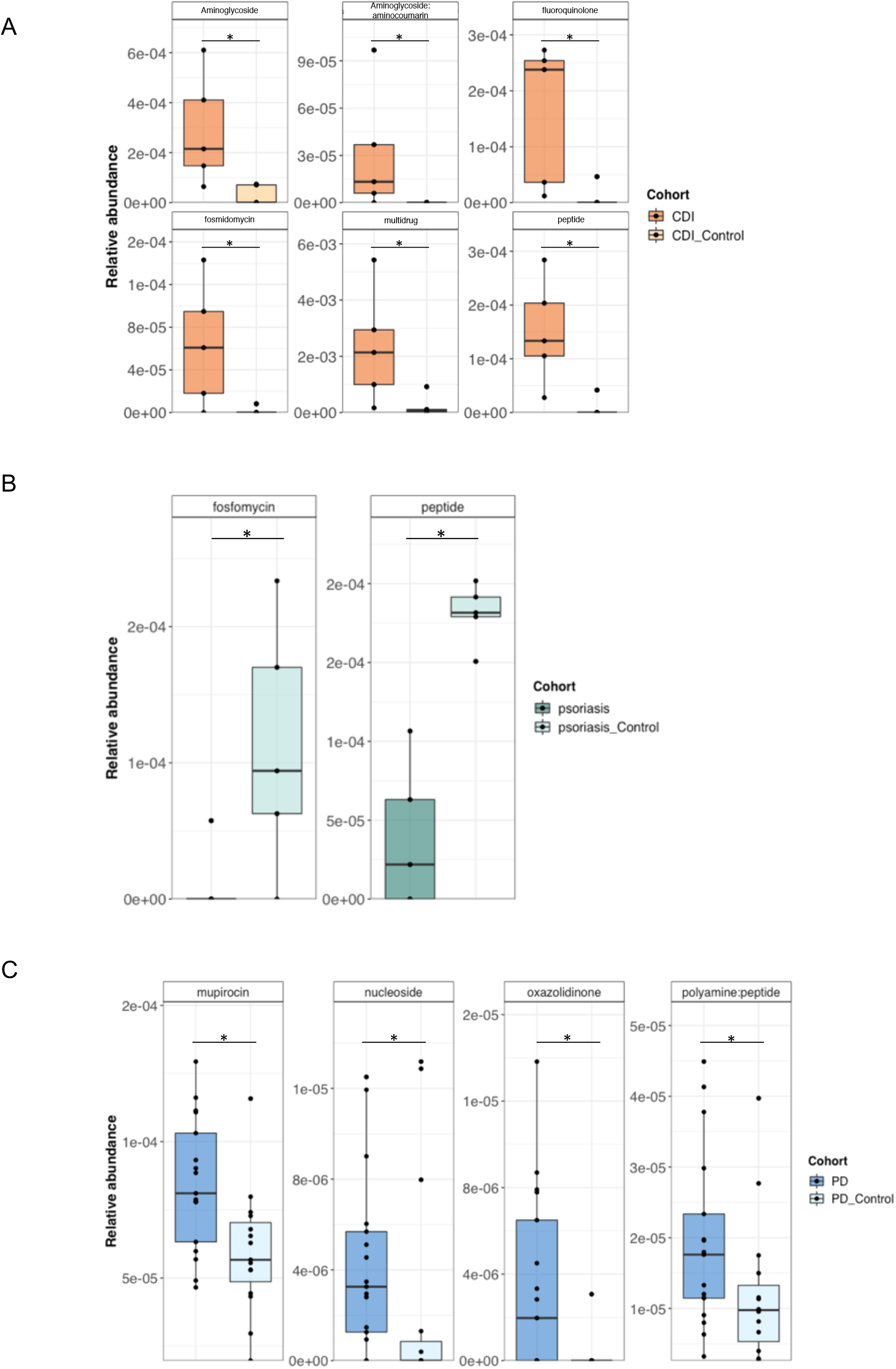
Antimicrobial resistance in three case-control metagenomic datasets. The relative abundance (%) of antimicrobial resistance categories statistically significantly differential abundant in; **a.** *Clostridioides difficile* infection versus control. **b.** psoriasis versus control. **c.** Parkinson’s disease vs control.* P-value < 0.05.

### Prediction of mobile genetic elements linked to virulence factors

Using the predictions generated by PathoFact, we resolved the genomic contexts and identified MGEs in all three case-control metagenomic datasets (Fig. 5a). Within all three datasets the presence of both phage- and plasmid-derived sequences was detected, although no significant difference was observed between diseased and control. Using PathoFact, we found that virulence factors, including bacterial toxins, contribute to the majority (~80%) of genes linked to MGEs in all datasets, with AMR contributing to the remaining MGEs (Fig. 5b). Furthermore, a small number of MGEs were found to be both linked to virulence factors as well as AMR.

**Figure 5.**
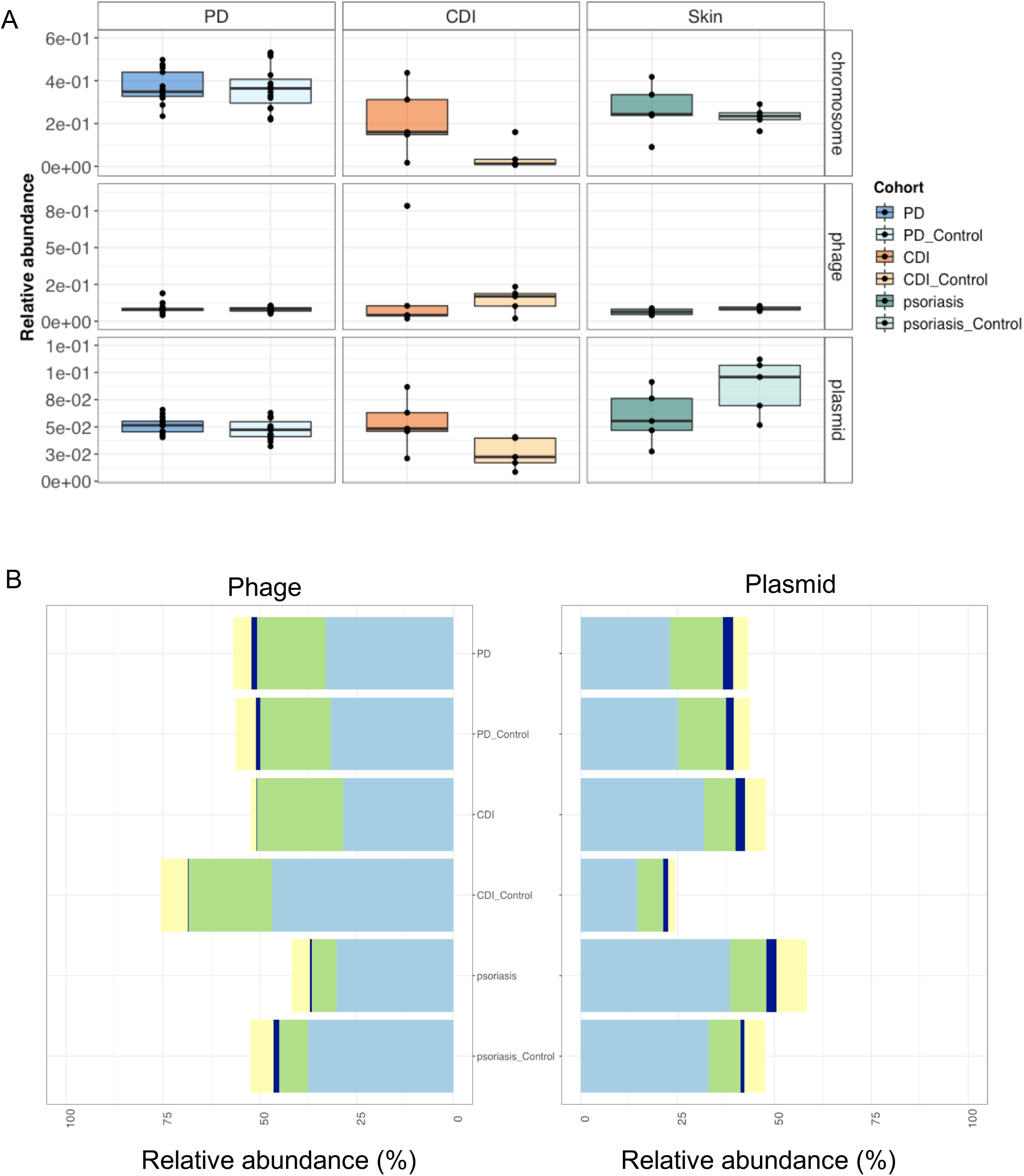
Identification of MGEs within three case-control metagenomic datasets. Relative abundance of MGEs within three metagenomic datasets *(Clostridioides difficile* infection, psoriasis (Skin) and PD). **a.** The overall relative abundance of phage and plasmids within the *Clostridioides difficile* infection, psoriasis and Parkinson’s disease datasets. **b.** The distribution of virulence factors, incl. toxins, and AMR between phage and plasmid in all datasets.

Of the ARGs linked to MGEs, the prevalence of the different resistance categories where identified using the pipeline. Within the CDI dataset the majority of the MGEs were linked to betalactam, MLS and multidrug resistance in both diseased and control groups (Fig. 6a). Additionally, a majority of both plasmids and phages were found to be linked to phenicol resistance within the control group. Within the skin metagenomes the majority of the resistance genes were linked to MGEs included beta-lactam and multidrug resistance in both diseased and control groups (Fig. 6b). Finally, of the resistance genes within the PD study both glycopeptide and multidrug resistance in both diseased and control groups were found to be linked primarily to both phage and plasmids (Fig. 6c).

**Figure 6.**
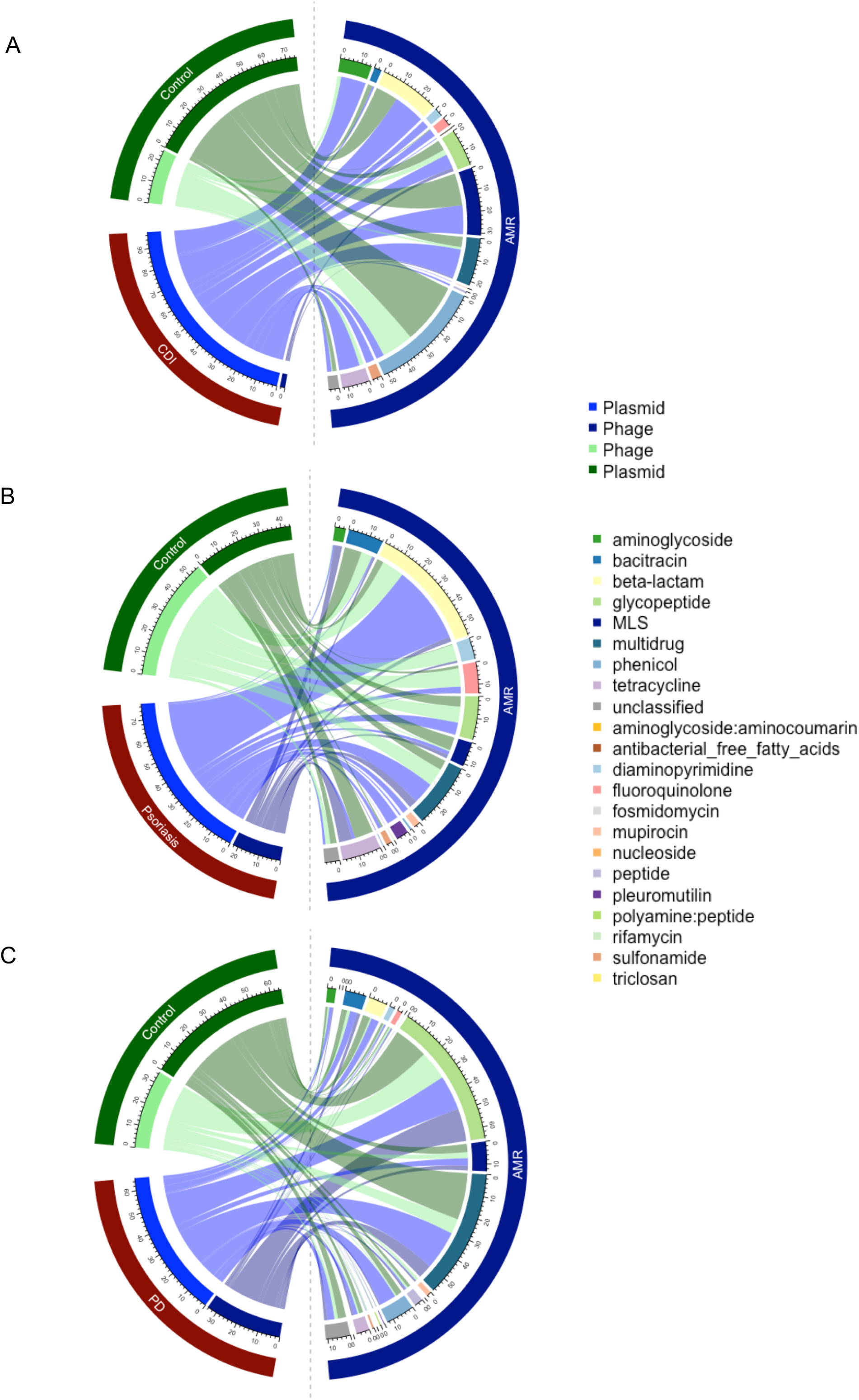
The prevalence of different resistance categories within the MGEs. **a.** prevalence of antimicrobial resistance genes within MGEs in *Clostridioides difficile* infection and control. **b.** psoriasis and control **c.** Parkinson’s disease and control.

## Conclusions

The identification of virulence factors, toxins and antimicrobial resistance genes are of immediate importance for understanding the pathogenic state of microbiomes. Using PathoFact we were able to identify virulence factors and bacterial toxins within three publicly available case-control metagenomic datasets. Furthermore, we were able to identify bacterial toxins differentially abundant between diseased and control groups in all datasets. Additionally, antimicrobial resistance genes were identified in all datasets with a significant difference of certain resistance categories between diseased and control individuals. The inclusion of MGEs is of particular importance in understanding the possible transmission of MGE-born virulence factors. With PathoFact we identified MGEs in all three datasets and were able to link these to the corresponding virulence factors, toxins and antimicrobial resistance genes.

Until now, no single pipeline has existed which has combined these distinct aspects. Furthermore, although several prediction tools exist for AMR, of which DeepARG has been chosen for its accuracy to be included in our pipeline, no or limited tools were available for the prediction of toxins and virulence factors. PathoFact utilizes the wealth of currently available software (e.g. AMR and MGE predictions) as well as newly generated tools (e.g. virulence factors and toxins). Furthermore, PathoFact can conveniently integrate updates and newly developed prediction tools. In conclusion, PathoFact combines the strength of AMR predictions linked to MGE predictions and integrates this with the prediction of toxins and virulence factors. PathoFact is a versatile and reproducible pipeline by its ability to run either the complete workflow or each module on its own, giving the investigator flexibility in their analysis.

## Authors’ contributions

LdN, SL, AHB and PW designed this study. LdN with support of SL, CCL, PM, AHB created the application. LdN and PW wrote the manuscript, CCL, PM and AHB contributed to the review of the manuscript. All authors have read and approved the manuscript.

## Acknowledgments

This work was supported by the Luxembourg National Research Fund (FNR) under grants PRIDE/11823097 to LdN, CCL, PM, PW and CORE/BM/11333923 to PW, the Michael J. Fox Foundation under grant No. 14701 and the European Research Council (ERC-CoG 863664) to PW. We are grateful for the feedback and beta-testing by Susheel Bhanu Busi and Susana Martinez Arbas. We are thankful to Valentina Galata for graphical support.

---

Supplementary Figure 1.

Antimicrobial resistance in three case-control metagenomic datasets. Relative abundance (%) of all identified resistance categories **a.** 25 antimicrobial resistance categories within *Clostridioides difficile* infection **b.** 24 antimicrobial resistance categories within the skin metagenome (psoriasis) and **c.** 26 antimicrobial resistance categories within the Parkinson’s disease study.

Supplementary Figure 2.

Distribution of virulence factors, incl. toxins and AMR over unclassified and ambiguous (predicted to be both plasmid and phage or phage and chromosome)

Supplementary Table 1.

Comparison of virulence factor prediction with MP3. Evaluated performance of the virulence prediction model versus the MP3 prediction tool regarding sensitivity, specificity and accuracy.

Supplementary Table 2.

Toxin domains differential abundant in diseased versus control in **a.** *Clostridioides difficile* infection, **b.** psoriasis and **c.** Parkinson’s disease.

