## Supplementary materials for "PathoFact: A pipeline for the prediction of virulence factors and antimicrobial resistance genes in metagenomic data"

Supplementary figure 1

A

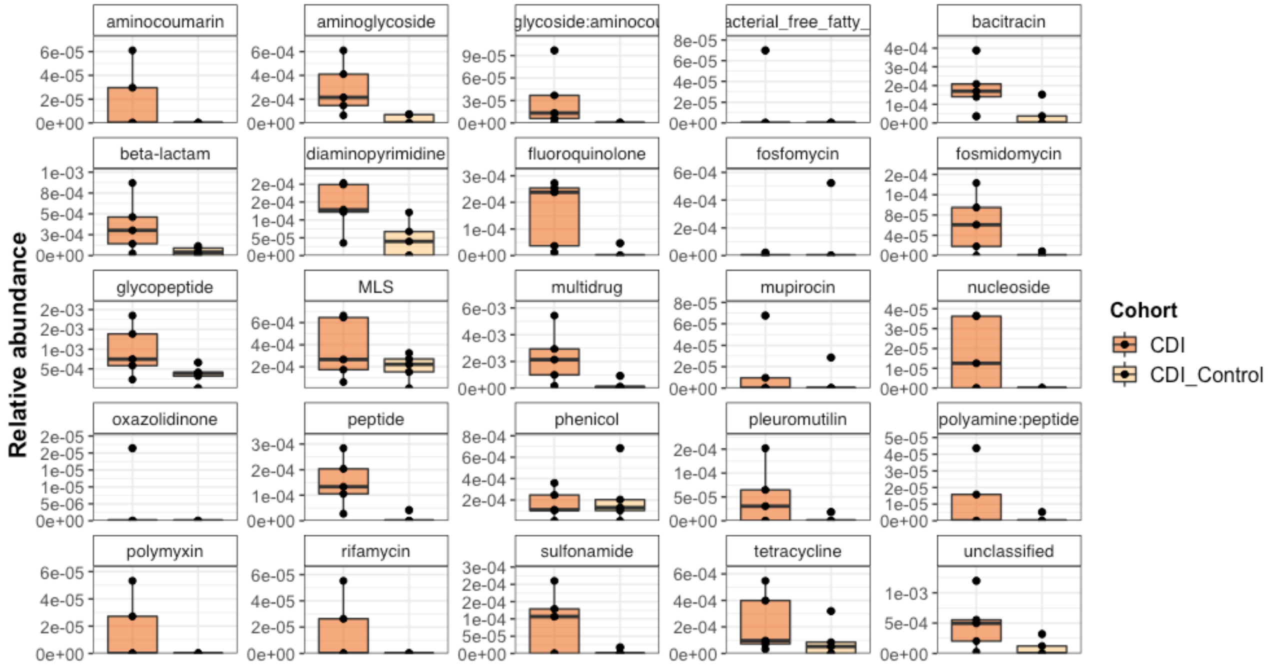


B

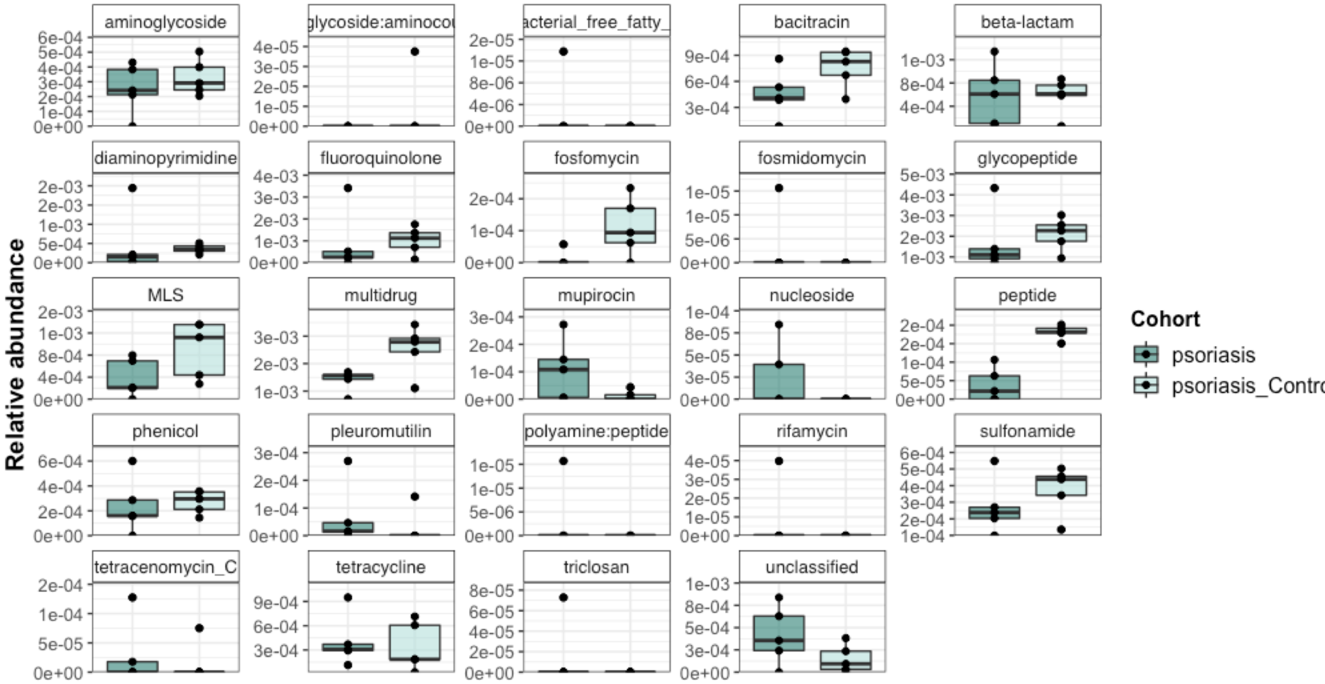

Supplementary figure 1

C

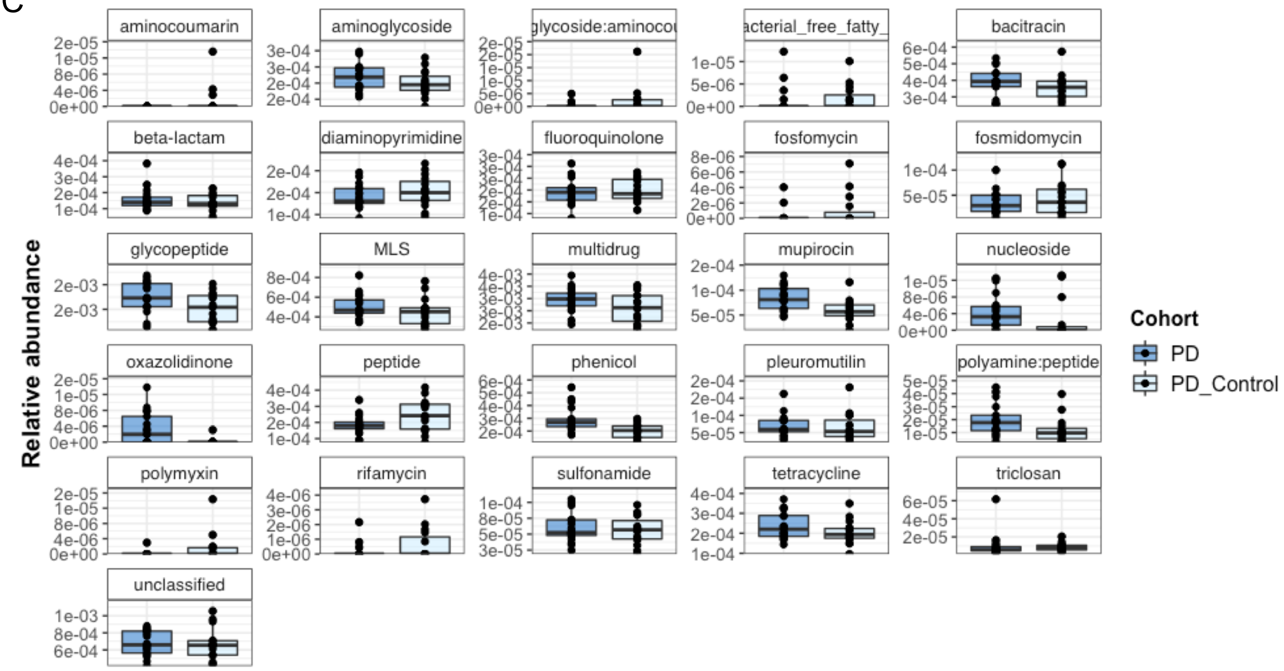

Supplementary figure 2

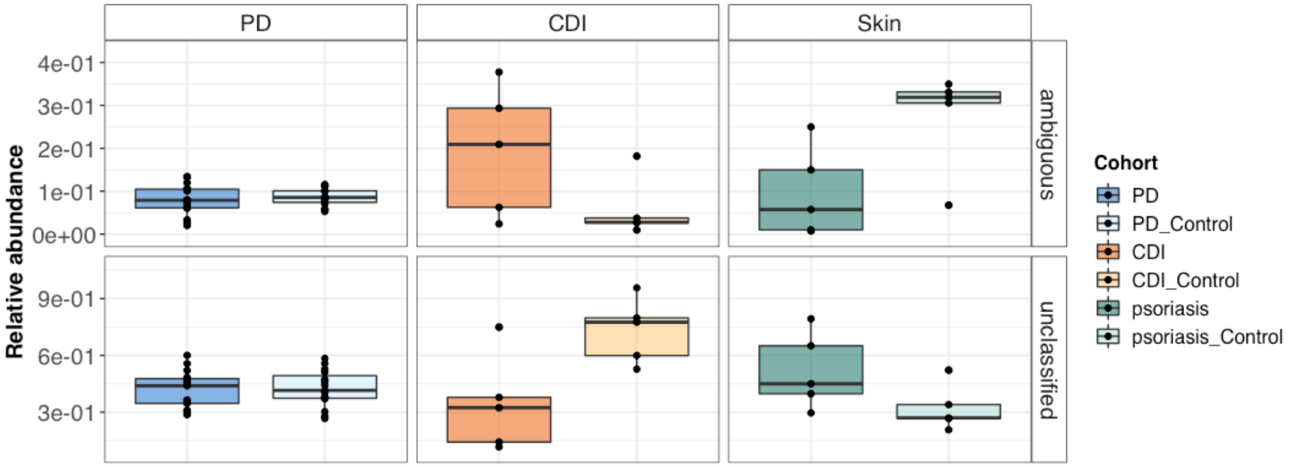


Supplementary Table 1

| | MP3 | Virulence factor prediction |
|---|---|---|
| Sensitivity | 0.125 | 0.886 |
| Specificity | 0.992 | 0.957 |
| Accuracy | 0.558 | 0.921 |

Supplementary Table 2

A

| HMM Domain | Log2FC | Name | Definition | Cohort |
|---|---|---|---|---|
| K11057 | -9,61 | cpb2 | Beta2-toxin | Control |
| K12788 | -6,20 | espH | LEE-encoded effector EspH | Control |
| K01387 | -5,94 | colA | Microbial collagenase | Control |
| K11023 | -5,01 | ptxA, artA | Pertussis toxin subunit 1 | Control |
| PF13945 | -4,59 | NST1 | Salt tolerance down-regulator | Control |
| PF08998 | -4,27 | Epsilon antitox | Bacterial epsilon antitoxin | Control |
| PF15534 | -3,83 | Ntox35 | Bacterial toxin 35 | Control |
| K11062 | -3,73 | entD | Probable enterotoxin D | Control |
| K11045 | -3,36 | cfa | cAMP factor | Control |
| PF15643 | -3,16 | Tox-PL-2 | Papain fold toxin 2 | Control |
| TIGR03396 | -3,10 | PC_PLC | Phospholipase C | Control |
| PF05015 | -2,13 | HigB-like toxin | RelE-like toxin of type II toxin-antitoxin system HigB | Control |
| K12340 | 3,02 | tolC | Outer membrane protein | Psoriasis |
| PF13935 | 4,70 | Ead/Ea22 | Ead/Ea22-like protein | Psoriasis |
| PF14449 | 4,78 | PT-TG | Pre-toxin TG | Psoriasis |
| K11052 | 5,20 | cylE | CylE protein | Psoriasis |

B

| HMM Domain | Log2FC | Name | Definition | Cohort |
|---|---|---|---|---|
| PF13954 | -5,84 | PapC_N | PapC N-terminal domain | CDI |
| PF06609 | -3,36 | TRI12 | Fungal trichothecene efflux pump | CDI |
| PF13953 | -2,90 | PapC_C | PapC C-terminal domain | CDI |

C

| HMM Domain | Log2FC | Name | Definition | Cohort |
|---|---|---|---|---|
| K10948 | -2.03 | hlyA | hemolysin | Control |
| PF15524 | -2.31 | Ntox17 | Novel toxin 17 | Control |
| PF09156 | 2.03 | Anthrax-tox_M | Anthrax toxin | PD |
| PF09599 | 2.18 | IpaC_SipC | Salmonella-Shigella invasion protein c | PD |
